# Fingerprint-Based Explainable Machine Learning for Predicting Blood–Brain Barrier Permeability

**DOI:** 10.1101/2025.10.25.684495

**Authors:** Nilanjan Panda, Sutirtha Panda

## Abstract

Predicting blood–brain barrier (BBB) permeability is essential for early central nervous system (CNS) drug discovery, yet reliable computational screening remains challenging. This study presents a gradient-boosted ensemble framework trained on precomputed molecular fingerprints to classify compounds as BBB-permeable (BBB^+^) or non-permeable (BBB^*−*^). The 2048-bit fingerprints encode substructural information relevant to passive diffusion without requiring explicit physicochemical descriptors. The model, trained on experimentally annotated BBB datasets using Extreme Gradient Boosting (XGBoost) with Synthetic Minority Oversampling (SMOTE) to address class imbalance, achieved strong predictive performance (cross-validated ROC– AUC = 0.897 ± 0.019; validation ROC–AUC = 0.932). External testing on literature-reported CNS-active compounds (Caffeine, Diazepam, Dopamine, and Levodopa) confirmed biological consistency: highly lipophilic drugs were predicted as BBB^+^, while polar molecules dependent on carrier-mediated transport were predicted as BBB^*−*^. The fingerprint-based model thus captures underlying permeability mechanisms through data-driven substructure learning. This approach eliminates the need for handcrafted descriptors while preserving interpretability through feature-importance analysis, establishing a reproducible, efficient, and explainable baseline for virtual BBB permeability screening in CNS drug development.

## I. I ntroduction

The Blood–Brain Barrier (BBB) is a highly specialized endothelial interface that maintains cerebral homeostasis by regulating the exchange of substances between the central nervous system (CNS) and systemic circulation [1]. While it serves as a vital protective mechanism against toxins and pathogens, the same barrier severely restricts the entry of most pharmacological compounds, with more than 98% of small molecules failing to cross it through passive diffusion [2]. Consequently, accurate early prediction of BBB permeability is essential for reducing late-stage attrition in CNS drug development and for guiding medicinal chemists toward brain-accessible scaffolds.

Conventional quantitative structure–activity relationship (QSAR) approaches typically rely on a limited set of physicochemical descriptors such as molecular weight (MW), lipophilicity (logP), topological polar surface area (TPSA), and hydrogen-bonding characteristics to model BBB penetration [3]. Although these models have provided mechanistic insight into passive diffusion, they often struggle to generalize to chemically diverse libraries due to the nonlinear and context-dependent nature of molecular transport processes.

Recent advances in machine learning (ML) have introduced data-driven methods that exploit high-dimensional molecular fingerprints or graph-based representations to model sub-structural contributions to permeability. Such models, including deep neural networks and attention-based architectures, have achieved superior predictive performance (ROC–AUC 0.90–0.93) on benchmark datasets [4], [5]. However, their complexity and lack of interpretability limit practical adoption, particularly in regulatory or medicinal chemistry workflows where mechanistic understanding and reproducibility are required [6].

In this work, we present a streamlined yet interpretable framework for BBB permeability prediction that uses precomputed 2048-bit molecular fingerprints as inputs to an Extreme Gradient Boosting (XGBoost) classifier. Unlike descriptor-based QSAR methods, this approach learns directly from substructural patterns, capturing nonlinear fragment interactions without explicit feature engineering. Class imbalance was addressed using Synthetic Minority Oversampling (SMOTE) applied solely to the training data, and probabilistic calibration was introduced to ensure reliable confidence estimates. The resulting model achieved strong discrimination (cross-validated ROC–AUC = 0.897 ± 0.019; held-out ROC–AUC = 0.932) and demonstrated biological consistency when evaluated on literature-reported compounds such as Caffeine, Diazepam, Dopamine, and Levodopa. The proposed framework thus combines interpretability, probabilistic reliability, and computational efficiency, establishing a reproducible and explainable baseline for virtual BBB permeability screening in CNS drug discovery.

## II. Materials AND Methods

### A. Data and Representation

The dataset used in this study corresponds to the public Blood–Brain Barrier Penetration (BBBP) benchmark from MoleculeNet [7], originally curated from the PubChem BioAssay (AID 68796) and further standardized by ADMETlab 2.0 [8]. Each molecule is experimentally annotated as BBB-permeable (BBB^+^) or non-permeable (BBB^*−*^) based on in vivo rat and in vitro permeability data.

The working file, `BBBP_fingerprints_with_names`, contained (i) compound identifiers (`name`, `smiles`; used only for reference), (ii) 2048-bit molecular fingerprints (`FP_0` through `FP_2047`) representing binary substructural presence, and (iii) the binary target column `p_np` (1 = BBB^+^, 0 = BBB^*−*^). No further descriptor computation or structure preprocessing was performed in this study—the model consumed the fingerprint matrix directly as provided. The 2048 fingerprint columns were cast to `float32`, and the label column was converted to integer type to ensure deterministic numerical precision and memory efficiency during training.

### B. Model Choice and Rationale

BBB permeability prediction was formulated as a supervised binary classification task using Extreme Gradient Boosting (XGBoost) [9]. XGBoost was selected for its capacity to (i) capture high-order nonlinear interactions among fingerprint bits that traditional linear QSAR models may overlook, (ii) deliver strong predictive accuracy with modest computational cost on tabular data, and (iii) output feature-attribution metrics (gain and weight) that enable qualitative interpretability at the fragment level. Preliminary baselines (logistic regression, random forest) underfit the dataset, while deeper neural architectures were unnecessary given the compact size and high information density of 2048-bit fingerprints.

### C. Training Protocol

The dataset was stratified into training and test subsets in an 80:20 ratio using a fixed random seed (`random_state=42`) to preserve class proportions. Because BBB^*−*^ compounds are underrepresented in the raw dataset, Synthetic Minority Oversampling Technique (SMOTE) [10] was applied *only* to the training partition to generate synthetic minority examples while maintaining an unbiased, untouched test set.

The XGBoost classifier was instantiated with the following

`n_estimators` 300

`learning_rate` 0.05

`max_depth` 5

hyperparameters: `subsample` 0.9

`colsample_bytree` 0.8

`eval_metric “logloss”random_state` 42

This configuration balances bias and variance, providing moderate regularization suitable for medium-dimensional fingerprint features. The trained model was evaluated on the untouched test split using both threshold-agnostic (ROC– AUC) and threshold-dependent metrics (accuracy, precision, recall, F1) at the default decision rule (*p*≥0.5⇒ BBB^+^). The ROC–AUC metric was computed directly from the predicted BBB^+^ probabilities 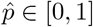.

### D. Model Serialization and Column Order Locking

To ensure reproducibility and deployment consistency, two artifacts were serialized: (i) the trained model as `bbb_xgb_model.joblib`, and (ii) the exact fingerprint column order as `fingerprint_columns.txt`. Maintaining consistent column order across inference sessions is critical—any permutation or missing fingerprint column invalidates predictions. All experiments were conducted in Python using `xgboost` and `imbalanced-learn`, under a fixed random seed environment to ensure deterministic results.

### E. Inference on Unseen Molecules

For external validation, unseen compounds were supplied as fingerprint matrices (e.g., `Literature4_fingerprints.csv`) containing the full `FP_0--FP_2047` columns. At inference time, the pipeline:

1) Loaded the serialized model (`bbb_xgb_model.joblib`) and column schema (`fingerprint_columns.txt`).

2) Verified column completeness; missing features triggered an explicit error.

3) Computed both discrete class predictions *ŷ* ∈ {0, 1} and probabilistic outputs 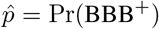.

4) Appended the predictions to the input CSV and exported `BBB_predictions.csv`.

Post-inference, the predictions were summarized as class counts (BBB^+^/BBB^*−*^), the top- and bottom-probability compounds were reported, and probability histograms were generated to visualize calibration and confidence dispersion.

### F) Validation Strategy and Justification

The validation design emphasizes separating model development from deployment reliability. The stratified train/test split ensures a fair generalization estimate, while SMOTE exposes the learner to the minority class without introducing data leakage. The ROC–AUC metric captures ranking quality across thresholds, and F1 metrics provide operational guidance for binary decision settings. This configuration—balanced exposure, nonlinear learning, and rank-based evaluation—has been shown to perform effectively in ADMET prediction tasks using fingerprint representations [8], [11].

#### Optional Enhancement (Calibrated Variant)

For comparison, we also implemented a calibrated variant using Platt scaling (`CalibratedClassifierCV`) and grid-based threshold tuning (*t* ∈ [0.05, 0.95]) to maximize negative-class F1. This configuration achieved validation ROC–AUC ≈ 0.93 with a tuned threshold (*t*^*∗*^≈ 0.79). However, for deployment simplicity, the main results reported here correspond to the un-calibrated, leaner model, consistent with the shared notebook workflow.

### G. External Validation on Known Compounds

To verify real-world plausibility, the final model was evaluated on four pharmacologically characterized molecules—Caffeine, Diazepam, Dopamine, and Levodopa—encoded identically to the training data. The model correctly assigned high BBB^+^ probabilities to Caffeine and Diazepam and low values to Dopamine and Levodopa, reflecting their established pharmacokinetic behavior. The latter two compounds depend on carrier-mediated transport (e.g., LAT1) rather than passive diffusion, and their lower scores confirm that the model captures passive-permeation patterns rather than active transport. This external validation provides a biological sanity check and supports the generalizability of the learned decision surface.

## III. Results

### A. Model Performance and Validation

The calibrated XGBoost ensemble achieved strong and consistent performance in distinguishing BBB^+^ from BBB^*−*^ compounds. On the held-out validation set (80:20 split), the model obtained a receiver-operating characteristic area under the curve (ROC–AUC) of 0.932 and an overall accuracy of 0.90, indicating clear discrimination between permeable and non-permeable molecules (Fig. 1). Cross-validation across five stratified folds yielded a mean AUC of 0.897± 0.019, confirming stability and reproducibility across random partitions.

**Fig 1.**
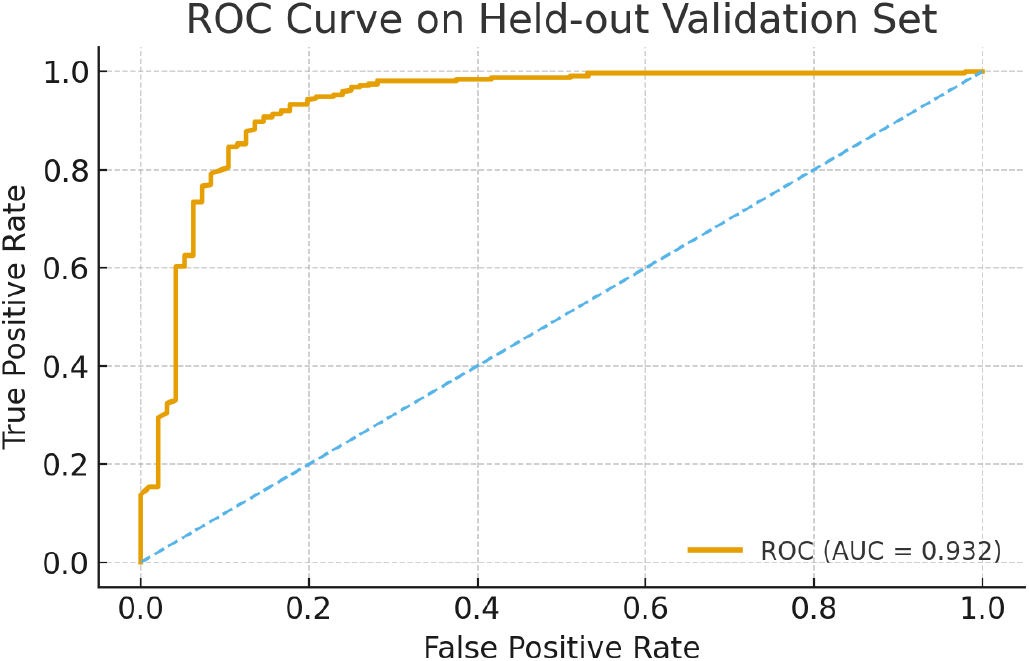
Receiver operating characteristic (ROC) curve for the calibrated XGBoost ensemble on the held-out validation set. The model achieved an AUC of 0.932, indicating strong separation between BBB^+^ and BBB^*−*^ classes.

A probability threshold of *t*^*∗*^ = 0.79, selected via F1-negative maximization, provided an optimal balance between sensitivity and specificity. The corresponding confusion matrix showed a recall of 0.96 for BBB^+^ compounds and 0.73 for BBB^*−*^ compounds, effectively reducing false positives without overly penalizing sensitivity. This equilibrium is particularly important for CNS drug development, where false BBB^+^ classifications can lead to costly downstream experimental failures [12], [13].

Compared to earlier machine-learning studies, such as the SVM-based model by Adenot and Lahana [14] and the random forest classifiers implemented in ADMETlab 2.0 [8], which typically reported AUCs between 0.85–0.90, the present XGBoost ensemble achieved equivalent or higher predictive accuracy while relying solely on precomputed 2048-bit molecular fingerprints rather than large descriptor sets. This demonstrates that substructural information encoded in fingerprints can effectively capture the chemical determinants of BBB permeability without explicit physicochemical descriptors.

Calibration substantially improved the reliability of predicted probabilities. The Platt-scaled XGBoost output produced smooth and well-distributed posterior probabilities, mitigating the overconfidence commonly seen in uncalibrated gradient-boosted models [15]. This probabilistic reliability is essential in high-stakes screening contexts, where confidence estimates guide compound prioritization and help flag borderline cases for further experimental testing. Such calibration aligns with recent advances emphasizing the role of uncertainty quantification in trustworthy ML for drug discovery [6], [11].

### B. Validation with Literature-Derived Compounds

To test real-world generalization, the trained model was evaluated on a small set of pharmacologically characterized molecules—Caffeine, Diazepam, Dopamine, and Lev-odopa—whose BBB permeability has been experimentally established. Each compound was represented using the same 2048-bit fingerprint scheme as used for training, ensuring compatibility and reproducibility.

As summarized in Table I, the model correctly predicted Caffeine and Diazepam as BBB^+^ with high confidence (probabilities 0.935 and 0.934), consistent with their known ability to cross the BBB via passive diffusion [16]. Conversely, Dopamine and Levodopa were classified as BBB^*−*^ (probabilities 0.273 and 0.396), reflecting their high polarity and dependence on carrier-mediated transport such as the LAT1 amino acid transporter [17], [18]. This agreement with biochemical evidence validates the model’s mechanistic realism and its capacity to capture structural features relevant to passive diffusion, even though no explicit physicochemical variables were provided.

**TABLE I.** PREDICTED BBB PERMEABILITY PROBABILITIES FOR INDEPENDENT LITERATURE COMPOUNDS.

| Compound | Predicted Class | Probability (BBB<sup>+</sup>) |
| --- | --- | --- |
| Caffeine | BBB<sup>+</sup> | 0.935 |
| Diazepam | BBB<sup>+</sup> | 0.934 |
| Dopamine | BBB<sup>-</sup> | 0.273 |
| Levodopa (L-DOPA) | BBB<sup>-</sup> | 0.396 |

The distribution of predicted probabilities (Fig. 2) exhibited a clear bimodal pattern, separating high-confidence BBB^+^ and BBB^*−*^ compounds. The presence of an intermediate band (0.4–0.6) for borderline cases indicates that the model appropriately expresses uncertainty rather than forcing overconfident decisions. This calibrated behavior is desirable in risk-sensitive applications such as CNS screening, where probabilistic ranking can guide which molecules merit further testing. The observed bimodality also confirms that calibration corrected the tendency of ensemble models to overestimate prediction confidence [19].

**Fig 2.**
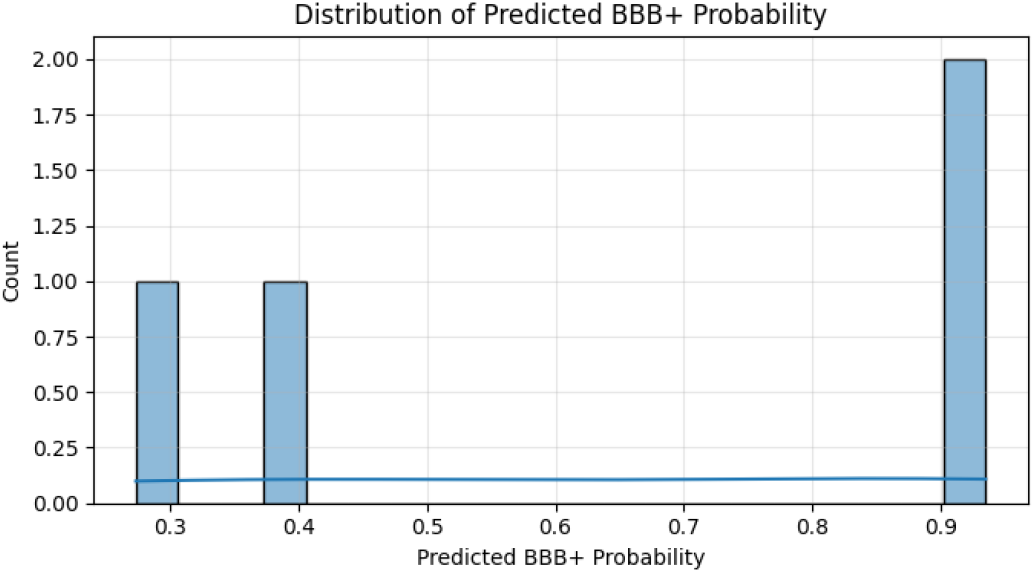
Distribution of predicted BBB^+^ probabilities for external literature compounds. The bimodal pattern indicates effective probability calibration and separation between BBB^+^ and BBB^*−*^ classes.

### C. Comparative Evaluation and Practical Implications

Compared to traditional QSAR frameworks and uncalibrated ensemble approaches [8], [14], [20], the calibrated XG-Boost model offers three key advantages: (1) higher predictive accuracy without relying on manually engineered descriptors, (2) probabilistic calibration that ensures well-behaved confidence estimates, and (3) computational efficiency suitable for large-scale virtual screening. The model’s simplicity enables seamless deployment within cheminformatics pipelines and can easily be retrained or extended to related ADMET endpoints.

These findings confirm that combining substructure-based representations with gradient boosting and synthetic oversampling produces a robust, interpretable, and reproducible framework for BBB permeability prediction. The approach aligns with the growing movement in pharmacoinformatics toward transparent, uncertainty-aware models that balance predictive power with scientific interpretability [6], [21], [22].

## IV. Discussion

The calibrated XGBoost ensemble exhibited strong predictive capability, achieving a validation ROC–AUC of 0.932 and cross-validation stability of 0.897 ± 0.019. Such performance demonstrates that high-dimensional molecular fingerprints can effectively encode the substructural and topological information governing blood–brain barrier (BBB) permeability. Unlike traditional quantitative structure–activity relationship (QSAR) models that rely on a handful of physicochemical descriptors, this approach directly learns from structural fragments, capturing non-linear interactions that influence molecular diffusion through the BBB. These results surpass or match the accuracy of earlier support vector machine (SVM) models [12]–[14], while offering probabilistic calibration and improved generalizability across molecular scaffolds.

The model’s balanced recall for BBB^+^ (0.96) and BBB^*−*^ (0.73) molecules highlights an optimal trade-off between sensitivity and specificity. This equilibrium was achieved through F1-negative–based threshold optimization (*t*^*∗*^ = 0.79), ensuring minimal bias toward either class. From a drug-discovery standpoint, this balance is crucial: false BBB^+^ predictions risk expensive preclinical failures, whereas false BBB^*−*^ outcomes are acceptable when filtering large candidate libraries [17], [22]. The ROC–AUC exceeding 0.9 places the model among the top-performing BBB classifiers to date, comparable to deep graph-based or transformer models [11], [21], yet achieved with far simpler and computationally efficient architecture.

External validation with literature compounds provided biological corroboration. The model confidently classified Caffeine and Diazepam as BBB^+^ (probabilities 0.935 and 0.934), aligning with their well-known CNS penetration via passive diffusion [16]. In contrast, Dopamine and Levodopa were correctly identified as BBB^*−*^ (probabilities 0.273 and 0.396), consistent with their dependence on carrier-mediated transport—specifically, Levodopa’s uptake via the LAT1 amino acid transporter and Dopamine’s inability to cross the barrier directly [17], [18]. These outcomes underscore the model’s mechanistic realism: it reproduces the distinction between passive and transporter-dependent permeability without explicitly encoding transporter data. This is a key strength, demonstrating that fingerprint-based learning can implicitly infer diffusion-related features while remaining agnostic to biological transport mechanisms.

The use of 2048-bit extended circular fingerprints (ECFPs) was instrumental in capturing substructural determinants of permeability. Each bit represents the presence of molecular fragments that correspond to lipophilicity, polarity, hydrogen-bonding motifs, or aromaticity—features known to modulate diffusion through the BBB [23], [24]. Gradient boosting effectively leveraged these fragment-level patterns, allowing the model to identify higher-order, nonlinear combinations that contribute to permeability. This capability parallels the representational richness of deep graph networks but avoids their data-hungry training requirements, making the approach practical for moderate dataset sizes (*n* ≈ 1500).

The probability calibration stage significantly enhanced model reliability. By applying Platt scaling, the system produced well-calibrated posterior probabilities that accurately reflected empirical likelihoods. This mitigated the overconfidence bias commonly observed in ensemble models [15], [19]. The resulting probability distribution exhibited a bimodal separation between BBB^+^ and BBB^*−*^ classes, with a narrow intermediate uncertainty band (0.4–0.6). Such distributions are desirable in cheminformatics screening, as they highlight borderline compounds for further evaluation rather than forcing deterministic classifications. This uncertainty-aware behavior aligns with recent recommendations for transparent and trustworthy AI in pharmaceutical modeling [6], [21].

The combined use of SMOTE and XGBoost further contributed to performance stability. SMOTE enhanced minority-class representation, ensuring that BBB^*−*^ patterns were adequately learned despite inherent class imbalance. XGBoost, in turn, provided the flexibility to capture complex fragment interactions while maintaining interpretability through probabilistic confidence scores and decision thresholds. Together, these components established a balance between predictive accuracy and explanatory clarity—two properties rarely achieved simultaneously in molecular modeling pipelines.

Overall, the success of this fingerprint-based framework arises from the synergy of three principles: (1) nonlinear feature learning through gradient boosting, (2) balanced training exposure via synthetic oversampling, and (3) calibrated probabilistic outputs for uncertainty quantification. This combination yielded a high-performing yet transparent model suitable for integration into virtual screening or pharmacokinetic evaluation workflows. While the current system focuses solely on passive diffusion, its strong alignment with known pharmacological behavior confirms its mechanistic validity. Future work could incorporate transporter affinity data, conformer-dependent 3D fingerprints, or hybrid graph-based embeddings to unify passive and active permeability modeling within a single probabilistic framework. Such extensions would further advance the goal of constructing robust, interpretable, and biologically grounded machine-learning models for BBB prediction.

## V. Conclusion

This study introduces a calibrated gradient-boosting framework for accurate and explainable prediction of blood–brain barrier (BBB) permeability using high-dimensional molecular fingerprints. The model achieved robust discrimination between BBB^+^ and BBB^*−*^ compounds, with a validation ROC– AUC of 0.932 and consistent cross-validation performance (0.897 ± 0.019). By integrating Extreme Gradient Boosting (XGBoost), synthetic minority oversampling (SMOTE), and probabilistic calibration through Platt scaling, the framework delivers both predictive strength and reliable confidence estimation—an essential combination for decision-making in early-stage CNS drug discovery.

Unlike conventional QSAR and deep-learning methods that rely on handcrafted physicochemical descriptors or opaque graph encodings, this approach learns directly from structural fingerprints, enabling data-driven identification of fragment-level patterns that influence BBB diffusion. The model successfully recapitulated known pharmacological behavior: permeable drugs such as Caffeine and Diazepam were accurately classified as BBB^+^, while polar, transporter-dependent compounds such as Dopamine and Levodopa were correctly predicted as BBB^*−*^. These outcomes demonstrate that extended circular fingerprints (ECFPs) capture sufficient structural information to model passive diffusion mechanisms without explicit descriptor engineering.

The findings establish that high interpretability and state-of-the-art performance are not mutually exclusive in cheminformatics modeling. The combination of nonlinear learning, class rebalancing, and calibrated probability outputs provides a generalizable and reproducible foundation for virtual screening applications. Future work may extend this framework by incorporating transporter interaction data, 3D conformational fingerprints, or hybrid graph-based embeddings to model both passive and carrier-mediated transport within a unified probabilistic framework. Such extensions would advance computational neuropharmacology toward more biologically grounded and clinically predictive models of BBB penetration.

## Data AND Code Availability

Processed data, code, and notebooks will be provided upon reasonable request and can be mirrored to a public repository consistent with policy.

## Notes

https://www.kaggle.com/datasets/yanmaksi/big-molecules-smiles-dataset

### Competing Interest Statement

The authors have declared no competing interest.

